# RodZ modulates geometric localization of the bacterial actin MreB to regulate cell shape

**DOI:** 10.1101/229286

**Authors:** Alexandre Colavin, Handuo Shi, Kerwyn Casey Huang

**Author notes:** These authors contributed equally (Alexandre Colavin, Handuo Shi). Corresponding author:Kerwyn Casey Huang,Stanford University, Department of Bioengineering,443 Via Ortega Shriram Center, Room 007,Stanford, CA 94305, USA.

## Abstract

In the rod-shaped bacterium *Escherichia coli*, the actin-like protein MreB localizes in a curvature-dependent manner and spatially coordinates cell-wall insertion to maintain cell shape across changing environments, although the molecular mechanism by which cell width is regulated remains unknown. Here, we demonstrate that the bitopic membrane protein RodZ regulates the biophysical properties of MreB and alters the spatial organization of *E. coli* cell-wall growth. The relative expression levels of MreB and RodZ changed in a manner commensurate with variations in growth rate and cell width. We carried out single-cell analyses to determine that RodZ systematically alters the curvature-based localization of MreB and cell width in a manner dependent on the concentration of RodZ. Finally, we identified MreB mutants that we predict using molecular dynamics simulations to alter the bending properties of MreB filaments at the molecular scale similar to RodZ binding, and showed that these mutants rescued rod-like shape in the absence of RodZ alone or in combination with wild-type MreB. Together, our results show that *E. coli* controls its shape and dimensions by differentially regulating RodZ and MreB to alter the patterning of cell-wall insertion, highlighting the rich regulatory landscape of cytoskeletal molecular biophysics.

## Introduction

Bacterial shape is determined by the cell wall, a cross-linked sugar network that is constantly remodeled as cells grow [1, 2]. In several rod-shaped organisms, cell-wall insertion is controlled by the cytoskeletal protein MreB [3, 4], a structural homolog of eukaryotic actin [5]. In *E. coli*, MreB forms oligomers [6] that bind the inner surface of the cytoplasmic membrane [7], rotate around the cell’s long axis in a manner that is dependent on activity of the essential cell-wall synthesis enzyme PBP2 [6, 8], and control the spatiotemporal pattern of cell-wall insertion [5, 9-11]. Disruption of MreB through point mutations [12-15], depletion [16], overexpression [16, 17], or antibiotics [16, 18, 19] can lead to subtle size changes or aberrant morphological phenotypes. Quantification of the pattern of MreB fluorescence as a function of geometry in exponentially growing cells [11] or in cell wall-deficient spheroplasts [20] revealed that MreB preferentially localizes to invaginations of the cell surface. Molecular dynamics simulations predicted that MreB polymers have nucleotide-dependent intrinsic curvature and substantial resistance to bending [21], both of which are key ingredients for sensing curvature. Moreover, simulations based on a mechanochemical model of cell-wall growth demonstrated that preferential localization to regions of negative Gaussian curvature is sufficient to straighten a bent cell [11]. Thus, biophysical feedback between cell shape and MreB-mediated wall growth appears to be crucial for cell-shape maintenance, although it remains unknown whether *E. coli* cells actively regulate the biophysical properties of MreB polymers to adjust cell shape and size.

*E. coli* cell shape has long been recognized to vary across growth phases, with cells becoming shorter as population optical density increases past ~0.3; cells become nearly round in stationary phase [22]. Moreover, the steady-state cellular dimensions of many rod-shaped bacteria adjust in response to nutrient-determined changes in growth rate [23, 24], with faster-growing cells having increased volume. The molecular mechanisms underlying changes in length and width are only partially understood, and there may be several pathways that indirectly affect cell size [24-26]. Nonetheless, mutation of a single residue of MreB to various amino acids was sufficient to drive a wide range of cell-size changes and to increase competitive fitness via decreases in lag time [14], suggesting that modification of MreB is a robust mechanism for determining cellular dimensions and thereby altering cellular physiology. Chemical inhibition of MreB polymerization by sublethal levels of the small molecule A22 resulted in dose-dependent changes to cell width and the chirality of cell-wall architecture [3], indicating that MreB polymeric properties may be biophysical parameters that can be exploited by the cell as tuning knobs for regulating cell width. How the geometric sensing function of MreB – which we define as MreB localization in response to morphological features such as surface curvature – is connected with cell size has not been systematically investigated. To elucidate the precise relationship between the molecular biophysics of the MreB cytoskeleton and the diverse landscape of cell shape requires both molecular structural investigations and precise single-cell experiments.

Here, we establish that the spatial organization of MreB in *E. coli* changes systematically across phases of growth, suggesting that the biophysical properties of MreB filaments alter in a manner commensurate with the nutrient-regulated changes in growth rate. Using single-cell microscopy, we determined that the protein RodZ [17, 27, 28] regulates the geometric sensing of MreB. Molecular dynamics simulations prompted us to propose that RodZ binding directly alters the conformational dynamics and intrinsic curvature of MreB polymers. We studied several MreB mutations that complement rod-like shape in the absence of RodZ when expressed alone or in combination with wild-type MreB (MreB^WT^). These mutants display enrichment of MreB to curvatures distinct from wild-type cells, and result in longer polymers. Simulations predict that these MreB mutations alter polymer bending dynamics in a manner consistent with the behavior of wild-type MreB bound to RodZ. Together, our findings demonstrate that regulation of RodZ tunes the geometric localization of MreB and thereby alters cell shape.

## Results

### *E. coli* cells rapidly change size as nutrients are depleted

Based on previous reports that *E. coli* cell mass decreases dramatically as the population increases beyond an optical density of ~0.3 [22], we hypothesized that passage through a typical growth curve would yield insights into the mechanisms of cell-size determination across a range of cell sizes in a single genotypic background. We interrogated a strain expressing the *mreBCD* operon under control of the native promoter on a plasmid, with a sandwich fusion of MreB to monomeric superfolder GFP (msfGFP) [11]. To monitor cell shape as a function of cell density, we back-diluted a 24-h, stationary-phase culture grown in lysogeny broth (LB) 1:200 into fresh LB in a test tube. Every 15 min, we extracted a small sample and imaged cells on an agarose pad with phase-contrast and epifluorescence microscopy to measure cell shape and MreB localization (Methods). We extracted cell contours [11, 14] and computed the mean length and width of the population at each time point (Methods). Concurrently, we measured optical density from a parallel culture to quantify bulk growth rate (Fig. 1a).

**Figure 1:**
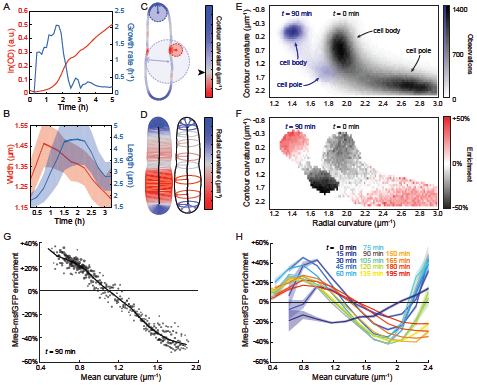
Cell shape and MreB localization patterns change as cell density increases in a growing culture. a) The population density and growth rate of *E. coli* cells growing in fresh LB were estimated from optical density (OD) measurements. b) Mean cell width and length across the population varied rapidly and asynchronously during the time course. Shaded areas represent standard deviation at each time point, whereas the variable thickness of the solid line represents the standard error at each time point (n > 800 cells). c-d) The local geometry of every point on each cell’s contour was characterized by the in-plane contour curvature (c) and perpendicular radial curvature (d). In (c), the red circle represents a point of negative contour curvature at the division site, the small blue circle represents a point of highly positive contour curvature at the poles, and the large blue circle represents a region of slightly positive contour curvature along the lateral wall. Black arrowhead next to colormap in (c) demarcates zero contour curvature, corresponding to flat regions. In (d), the radial curvature is inversely related to the local width of the cell. e) The frequency of pairs of contour and radial curvature values sampled by a population of *E. coli* cells after 0 min and 90 min of growth illustrates the range of curvature values across their surfaces. Each bin is 0.0821 um^-1^ (contour curvature) by 0.0165 um^-1^ (radial curvature). Black: *t* = 0, blue: *t* = 90 min. f) Enrichment of MreB fluorescence at *t* = 90 min observed for each bin in (e) with more than 50 observations demonstrates that MreB localization depends on both contour curvature and radial curvature. g) Each circle represents the MreB-msfGFP enrichment for the estimated mean curvature of a bin in (f) (corresponding to data from *t* = 90 min), which was defined as the average of contour and radial curvatures for that bin. The radius of each circle is linear with the log number of observations for the respective bin in (e). The weighted average across bins with a given mean curvature is shown as a solid line. h) The enrichment of MreB-msfGFP varied substantially across the growth curve. Shaded areas represent the standard deviation of enrichment from resampled data at each time point (Methods). All bins include at least 50 observations.

Along the growth curve of the cell culture’s exit from stationary phase, the mean cell length and width of the population rapidly changed. Within 1 h, there was a detectable increase in both mean width and length (Fig. 1b). Changes in width and length were not synchronized, with width initially increasing for the first 45 min, followed by a gradual decrease back to a value typical of stationary-phase cultures (~1 μm) over the growth curve (Fig. 1b). In contrast, length continuously increased for the first 1.5 h, plateaued for 45 min, and then gradually decreased (Fig. 1b). The cell population had not reached stationary-phase dimensions after 3 h, as the culture was still growing at a slow rate (Fig. 1a). Similar changes to cellular dimensions occurred in wild-type (unlabeled MreBCD at the native chromosomal locus) *E. coli* MG1655 cells (Fig. S1). Thus, cellular dimensions change in an asynchronous, nontrivial manner as nutrients are consumed, providing the opportunity to reveal connected changes in the behavior of the molecular mechanisms that construct the cell wall and determine shape.

### Curvature-based enrichment of MreB varies with cell density

To correlate MreB localization with features of a cell’s shape, we computed two curvature-related features at every point along the contour. Contour curvature describes bending along the cell outline (Fig. 1c). By our convention, the poles are regions of high positive contour curvature (small blue circle, Fig. 1c), while invaginations such as division-site constrictions are regions of high negative contour curvature (red circle, Fig. 1c). Although most of the cell away from the poles is approximately cylindrical, there are fluctuations in contour curvature (large blue circle, Fig. 1c) that we previously exploited to determine that MreB preferentially localizes to regions of negative contour curvature during exponential growth [11, 20]. The second curvature feature captures the local width, which is the distance of closest approach to the cell midline. We defined the inverse of this distance as the radial curvature (Fig. 1d), which approximates the out-of-plane curvature along the circumferential direction under the assumption of cylindrical symmetry. Thus, as cell width changes throughout the growth curve, straight regions have zero contour curvature regardless of cell width and smaller or larger radial curvature as cell width increases or decreases, respectively.

Throughout the growth curve after exit from stationary phase, cells adopted wide distributions of contour and radial curvatures (Fig. 1e). We computed the enrichment of MreB fluorescence as a function of contour and radial curvature across thousands of cells at each time point (Fig. S2). First, we calculated the distribution of curvatures as a two-dimensional histogram with fixed bin widths and bin positions for all cells in a given sample. Next, for each curvature bin, the intensity values of MreB localized at curvature values between the bin edges were averaged and normalized by the average fluorescence expected under the null hypothesis that MreB was randomly distributed along the contour of each cell. At 1.5 h, MreB was generally localized to negative contour curvature, as expected (Fig. 1f). However, the specific shape of the enrichment profile depended on the local radial curvature: MreB was more likely to be found at wider regions with negative contour curvature (Fig. 1f). Simulated microscopy [29] showed that this width-dependent enrichment could not be accounted for by optical artifacts due to the variable cell width (Fig. S3). Thus, for a fixed contour curvature, MreB prefers wider regions of the cell.

By contrast, at *t* = 0 (the beginning of the exit from stationary phase), MreB displayed a qualitatively distinct enrichment profile. Most notably, in the shorter and thinner stationary-phase cells (Fig. 1e, S2), MreB localized preferentially to the poles (high positive contour curvature) (Fig. 1f). Since the spatiotemporal patterns of MreB and of new cell-wall synthesis are highly correlated [11], this polar localization during stationary phase is consistent with our observation of rapid cell widening as cells exit stationary phase (Fig. 1b). To examine how the enrichment profile varied over time, we compressed the two-dimensional curvature into an approximate measure of the mean curvature (computed as the average of contour and radial curvature), binned mean curvatures into a onedimensional histogram, and recomputed the observation-weighted average of MreB enrichment as a function of mean curvature (Fig. 1g). To estimate the confidence of enrichment measurements, the enrichment profile was calculated 10 times from data bootstrapped from the original dataset with replacement, and the standard deviation of enrichment across resampled datasets was calculated for each bin. MreB localization was initially enriched in regions of high mean curvature (cell poles), but steadily decreased at later time points (Fig. 1h). Across the entire time course after the initial measurement (*t* = 0), there was a consistent enrichment of MreB at lower mean curvature (negative contour curvature). However, the enrichment profile varied quantitatively throughout the growth curve, with variations in enrichment on similar time scales as the changes in cellular dimensions (Fig. 1b). For example, cell width and length (Fig. 1b) were relatively constant between ~60 and 90 min, as was MreB enrichment (Fig. 1h). Moreover, MreB enrichment continued to change throughout the 3-h time course, as did cellular dimensions (Fig. 1b). Thus, MreB curvature sensing and cell shape both change dramatically as cells exit from stationary phase. Given the relationship between MreB and patterning of the insertion of cell wall material, we hypothesized that a direct relationship exists between curvature sensing and cell shape.

### Expression of RodZ alters MreB curvature enrichment

The large, systematic changes in MreB curvature-based localization suggest that molecular factors could be responsible for altering the subcellular behavior of MreB. Based on evidence from previous studies, a strong candidate for the regulation of MreB patterning is the bitopic membrane protein RodZ [17, 27, 28]. In *E. coli*, deletion of RodZ causes cells to become round [17, 28]. Changes to RodZ levels also tune cell shape: both underexpression and overexpression increase cell width, and overexpression also results in larger width variations [17]. Suppressor mutations of *rodZ* deletion that recover rod-like shape occur in *mreB* and *mrdA* (which encodes PBP2) [30]. Interestingly, most suppressors isolated in rich media die in minimal media, suggesting sensitivity to changes in a cellular quantity such as cell shape connected with growth rate [30]. Ribosomal profiling data indicated that there is approximately five-fold more MreB than RodZ in rich media, and that the ratio of MreB to RodZ abundance decreases in minimal media [31]. Similarly, mass spectrometry data from a variety of nutrient conditions showed that the ratio of MreB to RodZ generally decreases in nutrient-limited conditions [32]. Our mass spectrometry measurements of the strain used for our curvature enrichment measurements (Fig. 1) were consistent with these previous studies, and indicated that the MreB:RodZ ratio was ~30% higher in exponential phase than in stationary phase (Fig. 2a, Methods). Thus, we hypothesized that the MreB localization changes along the growth curve were driven by changes in RodZ expression relative to that of MreB.

**Figure 2:**
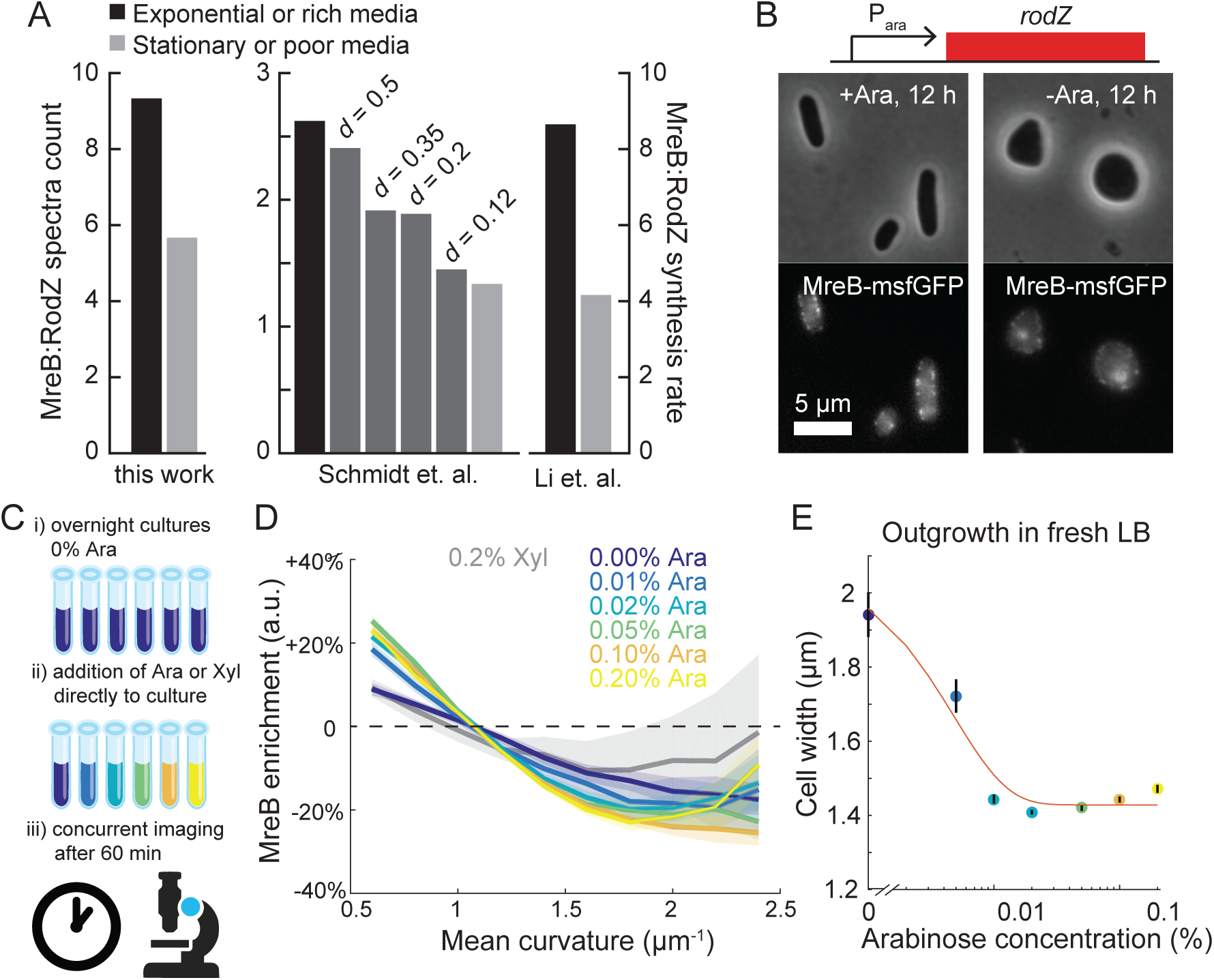
RodZ expression drives changes in MreB curvature enrichment profile. a) The ratio of MreB to RodZ protein abundance consistently increases in a manner concordant with growth rate across multiple independent studies. d: chemostat dilution rate. b) For a strain in which the native promoter of *rodZ* was replaced by P_ara_, RodZ expression is driven by arabinose (Ara). In the absence of arabinose, cells became spheroidal. c) Schematic of experimental approach in (d). Overnight cultures grown in the absence of arabinose were further incubated after adding varying amounts of arabinose. The distribution of MreB fluorescence was measured after 60 min. d) After 1 h of growth, induction of RodZ by arabinose enhanced the depletion of MreB at high contour curvature in a dose-dependent manner. By contrast, the enrichment profile was more uniform after induction with 0.2% xylose or 0% arabinose. Shaded areas represent the standard deviation of enrichment from resampled data at each condition. All bins include at least 50 observations. e) Overnight cultures grown in 0% arabinose and back-diluted 1:10,000 into fresh LB with varying levels of arabinose exhibited dose-dependent steady state widths after 4 h of growth. Black lines represent standard error of the mean (n > 50 cells).

To test how RodZ expression changes the curvature sensing of MreB and cell shape, we constructed a strain in which *rodZ* is deleted from the chromosome, and the native promoter of *rodZ* was replaced by P_ara_ (Methods). The strain background has a chromosomally integrated sandwich fusion of MreB to msfGFP as the sole copy of *mreB* [11]; the chromosomally integrated msfGFP fusion provides the best complementation of cell size of all MreB fusions studied to date [15]. As expected, after 12 h of growth in the absence of arabinose, cells were round (Fig. 2b), similar to Δ*rodZ* cells [17], whereas cells grown in the presence of arabinose were rod-shaped (Fig. 2b), albeit with larger cell widths than wildtype cells (Table S2).

To determine whether induction of RodZ changes MreB curvature sensing, we added various concentrations of arabinose (0 to 0.2%) directly to the 12-h culture of cells depleted of RodZ (0% arabinose), and imaged cells after 60 min (Fig. 2c). Since cells had already depleted the nutrients, little to no increase in optical density took place during the 60 min of arabinose treatment (Fig. S4a). For all cultures, there was enough cell-shape variability to measure mean curvature enrichment profiles of MreB fluorescence. The culture grown without arabinose had a relatively flat enrichment profile, signifying approximately random localization. As the arabinose concentration was increased, we measured increased enrichment of MreB to lower mean curvature and depletion at high curvature (Fig. 2d). Importantly, the increased range of enrichment with arabinose induction (-25% to 25%) was in reasonable quantitative agreement with the profiles we measured during wild-type outgrowth from stationary phase (Fig. 1h). Moreover, overnight cultures grown in the absence of arabinose and then back-diluted 1:10,000 in fresh LB with varying levels of arabinose exhibited a dose-dependent average width after 4 h, with higher arabinose concentrations leading to wildtype-like widths (Fig. 2e). By contrast, cells grown in xylose rather than arabinose exhibited MreB curvature enrichment similar to the original overnight culture (0% arabinose) (Fig. 2d). In addition, cells grown with xylose exhibited more diffuse MreB fluorescence than cells grown with arabinose, as measured by the difference in the distribution of peripheral fluorescence values across the population (Fig. S4b). These results show that RodZ expression alters MreB localization in a dose-dependent manner, driving enhanced curvature sensitivity, which further regulates cell shape.

### MreB mutants that suppress *ΔrodZ* growth defects have altered cell shape and curvature sensing

A previous study identified several mutations in MreB that were selected as suppressors of the slow-growth phenotype of Δ*rodZ* cells [30]. All of these mutants were able to grow as rods in the absence of *rodZ* [30]. We were interested to determine whether these mutations also modify the curvature sensing of MreB. We introduced three of these MreB mutations (D83E, R124C, and A174T) into *E. coli* MG1655 cells expressing the msfGFP sandwich fusion to MreB as the sole copy at the native *mreB* locus (Methods). These three mutations were selected to cover multiple regions of MreB; A174 is near the RodZ binding interface in domain IIA, R124C is near the membrane binding interface in subdomain IA, and D83E is at the double-protofilament interface in subdomain IB [33] (Fig. 3a). We also investigated a fusion of MreB^E276D^ to msfGFP, since E276 is located at the polymer interface and near the RodZ binding interface (Fig. 3a).

**Figure 3:**
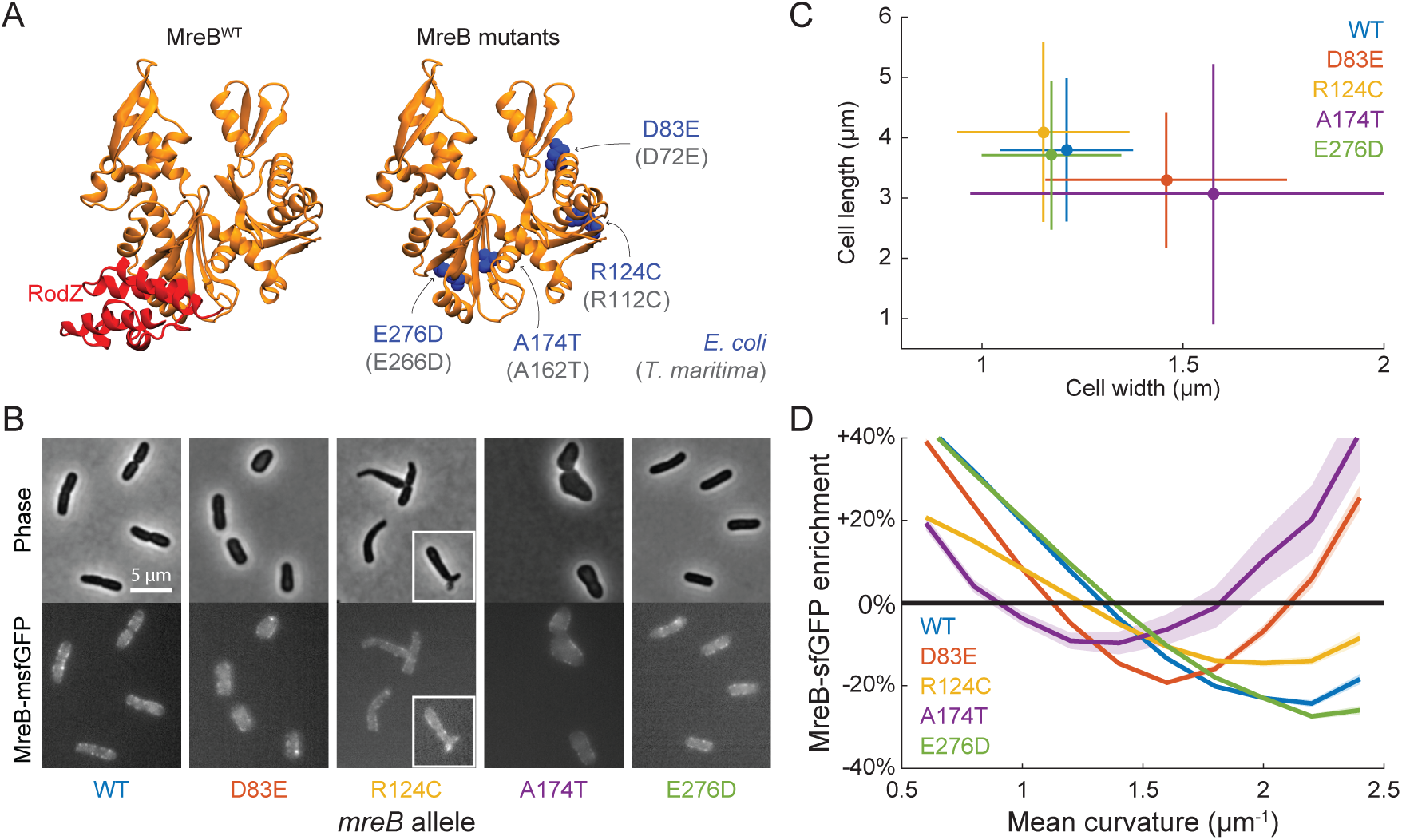
Genetic perturbations that alter the MreB curvature enrichment profile. a) RodZ binds near the polymerization interface in domain IIA of MreB (left). Mutations in MreB previously identified [30] to suppress Δ*rodZ* growth defects (D83E, R124C, and A174T), as well as a mutation at the polymerization interface (E276D), are spread throughout the protein and are conserved in *T. maritima* (gray text in parentheses; right). b) MreB mutants (all as sandwich fusions to msfGFP) have a variety of cell shapes, including wider cells (MreB^D83E^), tapered and sometimes branched cells (MreB^R124C^; white box highlights a branched cell), much wider and rounder cells (MreB^A174T^), and wild-type-like cells (MreB^E276D^). Error bars represent ±1 standard deviation (*n* > 230 cells). c, d) Strains that harbor mutations in MreB that suppress Δ*rodZ* growth defects have altered cellular dimensions (c) and MreB curvature enrichment profiles (d) relative to wildtype, whereas MreB^E276D^ cells are similar to wildtype. Error bars in (c) represent standard deviation of population. All bins in (d) include at least 50 observations.

MreB^D83E^ and MreB^R124C^ cells had a small, but significant, decrease in maximal growth rate compared to wild-type cells, while the maximal growth rate of MreB^A174T^ cells was almost two-fold lower than MreB^WT^ cells (Fig. S5a,b). These three strains also had significantly longer lag times than wild-type cells (Fig. S5c). We measured the cellular dimensions of each strain at the time of reaching maximal growth rate (Fig. 3b, c). MreB^D83E^ and MreB^A174T^ cells were somewhat and much wider and shorter than MreB^WT^ cells, respectively (Fig. 3b,c), while MreB^124C^ cells had a average width and length similar to wild-type cells (Fig. 3c) but exhibited substantial tapering and occasional branching (Fig. 3b). MreB^E276D^ cells had quantitatively similar growth (Fig. S5a) and shape (Fig. 3b,c) phenotypes to those of wild-type cells. These shape phenotypes are in good agreement with the study in which they were originally identified [30].

We next quantified the localization of MreB fluorescence as a function of curvature at the time when each strain reached its maximum growth rate. Since some of the strains exhibited more aberrant shapes than MreB^WT^ (MreB^R124C^ and MreB^A174T^ in particular), we normalized the MreB enrichment (Fig. S6a) by the enrichment calculated from the fluorescence signal from the membrane dye FM4-64 (Fig. S6b), and observed qualitatively similar results without (Fig. 3d,S6a) and with (Fig. S6c) normalization. MreB^E276D^ cells had a curvature enrichment profile that was very similar to that of MreB^WT^ (Fig. 3d). MreB^D83E^ and MreB^A174T^ cells had profiles shifted such that the crossover curvature at which localization was random (enrichment = 0) was lower for wider cells (Fig. 3d). MreB^R124C^ cells had a flatter enrichment profile than any of the other strains, indicating less curvature sensitivity; this finding suggests that the branching that we observed in MreB^R124C^ cells (Fig. 3b) is due to increased potential for growth at the polar (high curvature) regions during cell division, as has been observed in various cell-wall mutants [34]. Thus, while the three Δ*rodZ* suppressor mutants drive rod-shaped growth in the absence of RodZ, they have non-wild-type growth, shape, and localization phenotypes.

### Molecular dynamics (MD) simulations suggest that RodZ alters the intramolecular conformations and filament properties of MreB

Since the cytoplasmic tail of RodZ directly binds MreB [7], we hypothesized that the altered curvature sensing of MreB due to RodZ expression is driven by a direct biophysical interaction that alters the conformation of MreB filaments. We previously used all-atom MD simulations to demonstrate that *Thermotoga maritima* MreB subunits adopt a range of conformations connected with filament properties [21], and predicted a polymerization-induced flattening of MreB subunits that was subsequently verified using X-ray crystallography [33]. The conformations of MreB filaments can be described by the intermolecular bending (out-of-plane θ_1_, in-plane θ_2_) and twisting (θ_3_) between two adjacent MreB subunits (Fig. 4a,b). To investigate whether RodZ binding alters the conformations of MreB filaments, we used the *T. maritima* MreB-RodZ co-crystal structure (PDB ID: 2WUS) to simulate MreB dimers with RodZ bound to both subunits (Methods, Fig. 4c). In the absence of RodZ, an ATP-bound MreB dimer exhibited significant bending along θ_2_ compared to ADP-bound dimers (Fig. S7), as we previously reported [21], as well as some bending along θ_1_ (Fig. 4d). In contrast, in the presence of RodZ, dimer bending was drastically reduced along both the θ_1_ and θ_2_ bending axes (Fig. 4d, S7). These results were consistent across replicate simulations (Fig. S7). Thus, since RodZ directly alters the spectrum of conformations adopted by MreB dimers *in silico*, we predict that the binding of RodZ to a fraction of the MreB subunits, which will be related at least in part to the stoichiometry of MreB and RodZ (Fig. 2a), causes altered curvature sensing of MreB *in vivo*.

**Figure 4:**
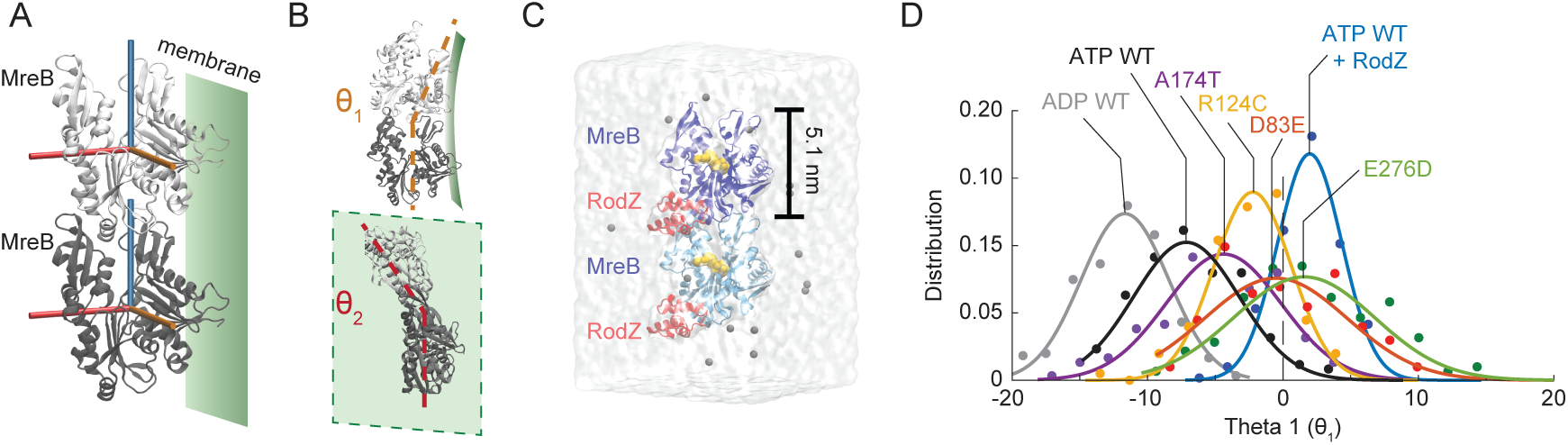
RodZ binding and MreB mutations may alter the bending properties of MreB filaments. a) Schematic of an MreB dimer (PDB ID: 1JCG) and its orientation relative to the membrane. The bending of MreB protofilaments is captured by the relative orientation along three orthogonal axes (cylinders) of adjacent MreB subunits, colored light (top) and dark gray (bottom). The membrane binding interface of the MreB protofilament is shown as a green plane. b) Schematic of MreB dimer bending angles out of the plane of the membrane (θ_1_, top) and in the plane of the membrane (θ_2_, bottom). In the crystal structures that form the initial states of our MD simulations, these bending angles are zero. c) MD simulation system comprised of a nucleotide-bound *T. maritima* MreB dimer, with each subunit bound to the cytoplasmic tail of RodZ. d) D72E, R112C, A162T, and E266D mutations in ATP-bound *T. maritima* MreB shift the θ_1_ bending angles toward that of a RodZ-bound ATP dimer, signifying filament straightening.

### Mutations in MreB also lead to straighter filaments, mimicking RodZ-bound MreB

Since RodZ expression modulates MreB curvature enrichment, and since our simulations predicted that RodZ-binding alters MreB filament mechanics, we asked whether the Δ*rodZ* suppressor mutants we studied *in vivo* also exhibit smaller bending angles than MreB^WT^, indicating straighter polymers. Since all three positions are conserved in *E. coli* and *T. maritima*, as is E276, we carried out all-atom MD simulations of dimers of the corresponding *T. maritima* mutants (D72E, R112C, A162T, E266D; Fig. 3a) bound to ATP, in the absence of RodZ. We observed shifts in the bending angles θ_1_ and θ_2_ toward zero (the approximate bending angle for MreB^WT^ bound to RodZ; Fig. 4d, S7) for all mutants, with a high degree of reproducibility in replicate simulations (Fig. S7). The MreB^A162T^(MreB^A174T^) mutant, which had a shape phenotype (Fig. 3b) closer to that of spherical RodZ-cells (Fig. 2b), displayed only a small degree of straightening, while the MreB^D72E^(MreB^D83E^), MreB^R112C^(MreB^R124C^), and MreB^E266D^(MreB^E276D^) dimers showed intermediate straightening. These data further support the hypothesis that MreB filament mechanics is an important component of cell-shape regulation.

### A combination of MreB^WT^ and MreB^E276D^ recovers rod-like shape in the absence of RodZ

In our MD simulations, MreB^E266D^(MreB^E276D^) dimers displayed intermediate straightening (Fig. S7), suggesting that the mechanical properties of polymers of this mutant are different from those of MreB^WT^ despite having similar growth and shape phenotypes to wildtype in the presence of RodZ (Fig. 3b-d, S5). Thus, we wondered whether MreB^E276D^ cells would be rod-shaped in the absence of RodZ. We constructed a Δ*rodZ* strain with MreB^E276D^ fused to sfGFP [15]. Like MreB^WT^ cells, MreB^E276D^ cells were round in the absence of RodZ (Fig. S5d) and grew as slowly as Δ*rodZ* MreB^WT^ cells, indicating that MreB^E276D^ is not a suppressor of Δ*rodZ* growth defects. Thus, we hypothesized that the mutant imitates a scenario in which MreB constitutively binds RodZ, similar to overexpression of RodZ, which is known to disrupt rod-like shape and result in round cells [17].

To test this hypothesis, we sought to create a genetic background in which only a fraction of MreB is bound to RodZ. We constructed a strain expressing MreB^WT^ from the chromosome and MreB^E276D^-msfGFP from a plasmid with the otherwise wild-type *mre* operon driven by the native *mreB* promoter (Methods) in a Δ*rodZ* background; we will refer to this strain as Δ*rodZ* MreB^WT^+MreB^E276D*^ (the asterisk indicates the presence of msfGFP), and use similar notation for strains with other *mreB* alleles on the chromosome and fused to sfGFP on the plasmid, with the rest of the *mre* operon included on the plasmid in all such strains. The Δ*rodZ* MreB^WT^+MreB^E276D*^ strain formed larger colonies and grew more quickly than Δ*rodZ* MreB^WT^ cells (Fig. S5). Δ*rodZ* MreB^WT^+MreB^E276D*^ cells were rod-shaped (Fig. 5a), albeit wider than MreB^WT^ cells (Fig. S8, Table S2), indicating that complementation was not perfect. To determine whether the MreB copy number is important for recovering of rod-shape, we constructed Δ*rodZ* MreB^WT^+MreB^WT*^ and Δ*rodZ* MreB^E276D^+MreB^E276D*^ strains. Δ*rodZ* MreB^WT^+MreB^WT*^ cells were slow-growing spheres (Fig. 5b, S5a,b), unlike Δ*rodZ* MreB^WT^+MreB^E276D*^ cells that displayed a contour curvature profile representative of rod-shaped cells (Fig. 5b). After 90 min of growth, the Δ*rodZ* MreB^WT^+MreB^E276D*^ strain exhibited a wider range of MreB enrichment than the Δ*rodZ* MreB^WT^+MreB^WT*^ strain (Fig. 5c). While some Δ*rodZ* MreB^E276D^+MreB^E276D*^ cells were rod-shaped (Table S2), this strain displayed significantly longer lag time (Fig. S5a,c) and lower growth rate (Fig. S5a,d) than Δ*rodZ* MreB^WT^+MreB^E276D*^ or Δ*rodZ* MreB^WT^ cells (Fig. S5a,c,d).

**Figure 5:**
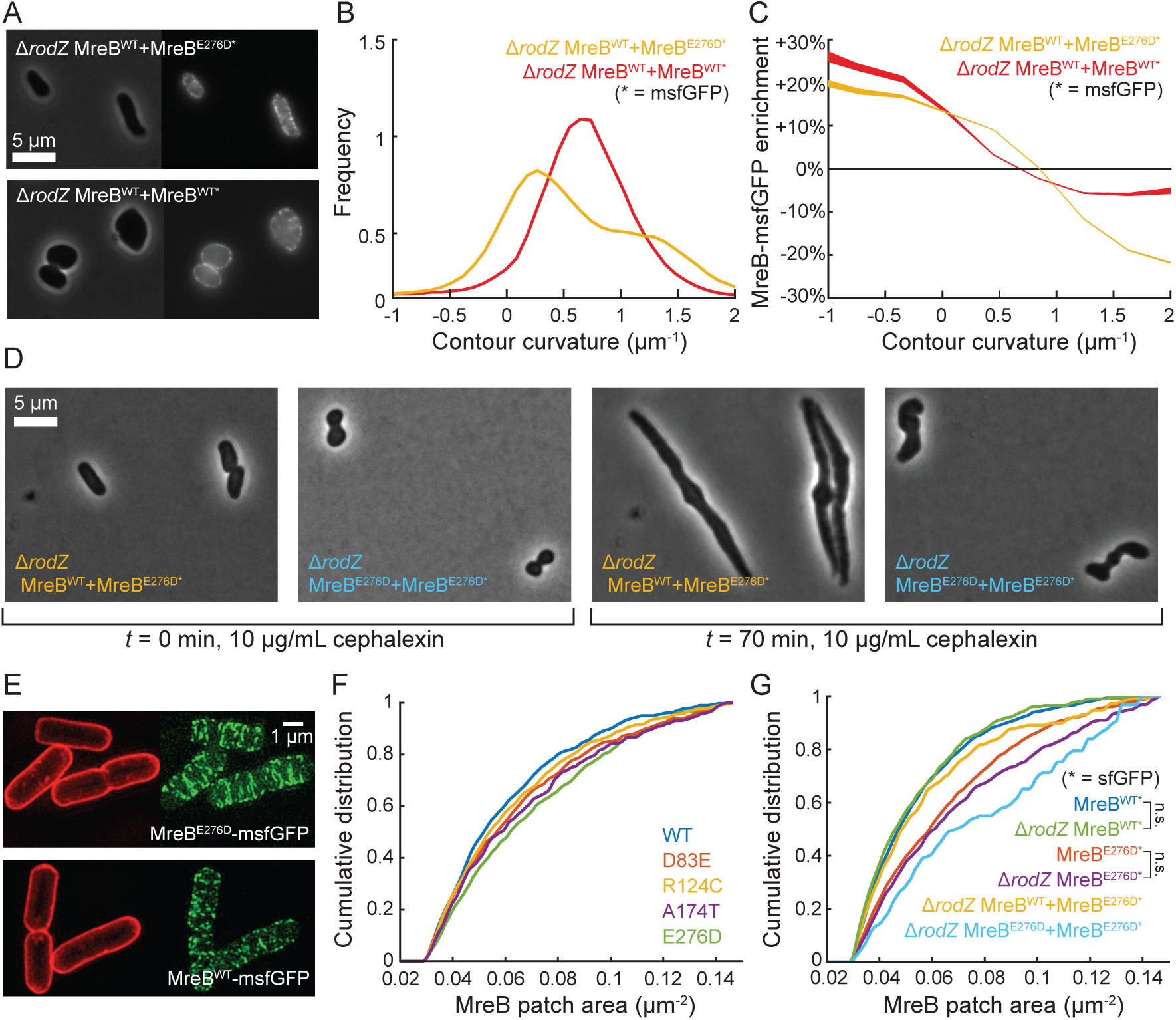
Concurrent expression of MreB^E276D^-msfGFP and MreB^WT^ recovers rod shape in Δ*rodZ* cells. a) Rod shape was rescued in Δ*rodZ* cells with chromosomal expression of MreB^WT^ by introducing a plasmid-borne copy of MreB^E276D^-msfGFP (top), but not with a plasmid-borne copy of MreB^WT^-msfGFP (bottom). b) Δ*rodZ* MreB^WT^+MreB^E276D*^ cells exhibited a bimodal contour curvature distribution with one peak centered near zero, characteristic of rod-shaped cells, unlike the unimodal distribution of Δ*rodZ* MreB^WT^+MreB^WT*^ cells. c) Δ*rodZ* MreB^WT^+MreB^E276D*^ cells exhibited enhanced depletion of MreB^E276D^-msfGFP at high contour curvature as compared with MreB^WT^-msfGFP in Δ*rodZ* MreB^WT^+MreB^WT*^ cells. d) Δ*rodZ* MreB^WT^+MreB^E276D*^ cells maintained a rod-like shape even when division was inhibited by cephalexin (10 μg/mL), while Δ*rodZ* MreB^E276D^+MreB^E276D*^ cells often failed to elongate by two-fold in 70 min (50% vs. 80% Δ*rodZ* MreB^WT^+MreB^E276D*^ cells) and lost the ability to regulate cell width. Snapshots of cells shown before (left) and after (right) 70 min of cephalexin treatment. e) Structural illumination microscopy revealed that the MreB^E276D^-msfGFP mutant strain had qualitatively longer filaments than the strain harboring MreB^WT^-msfGFP. Red fluorescence represents FM4-64FX membrane staining. f) The cumulative distributions of MreB-msfGFP fluorescence patch sizes for Δ*rodZ* suppressor MreB mutants were intermediate between those of MreB^WT^ and MreB^E276D^. Each MreB patch was defined as a continuous region larger than the diffraction limit with high GFP signal located within cell contours. g) Patch sizes for various strains varied widely, as indicated by the cumulative distribution of MreB-msfGFP fluorescence. Strains containing MreB^E276D^-msfGFP consistently showed distributions indicative of larger patch sizes (*p* < 10^-18^, *t*-test). Deletion of RodZ alone did not lead to statistically significant differences in the distribution of MreB patch sizes.

To ascertain whether Δ*rodZ* MreB^WT^+MreB^E276D*^ and Δ*rodZ* MreB^E276D^+MreB^E276D*^ cells were truly undergoing rod-like growth, we grew them in the presence of cephalexin, which inhibits the division-specific cell wallsynthesis enzyme PBP3 [35]. In the presence of cephalexin, round *E. coli* cells lyse due to the inability to divide [36], whereas rod-shaped *E. coli* cells elongate to tens of microns [11]. Many Δ*rodZ* MreB^WT^+MreB^E276D*^ cells (80%) elongated by more than two-fold over 70 min without dramatically changing cell width, whereas only 50% of Δ*rodZ* MreB^E276D^+MreB^E276D*^ cells were able to double in length as opposed to halting growth (Fig. 5d, *n* ≥ 50 cells). Taken together, these Δ*rodZ* MreB^WT^+MreB^E276D*^ data demonstrate recovery of rod-like shape in the absence of RodZ; the enhanced recovery compared with Δ*rodZ* MreB^E276D^+MreB^E276D*^ cells suggest the importance of the relative levels of RodZ-bound and unbound MreB in wild-type cells.

### RodZ-related MreB mutant filaments are longer than MreB^WT^ filaments

How are the biophysical properties of MreB connected to the cellular-scale properties of filaments and intracellular patterning? Since our MD simulations predicted that RodZ binding and various MreB mutations alter curvature enrichment by changing filament bending, and moreover since some perturbations (A174T, E276D) occurred near the polymerization interface of the MreB filament structure, we wondered whether these changes manifested in part as increased MreB polymer length. MreB usually forms diffraction-limited puncta [9, 10]; thus, to measure the patterning of MreB, we carried out super-resolution imaging using structured illumination microscopy (Methods). Strikingly, some cells expressing MreB^E276D^ alone contained filaments that were much longer than filaments in MreB^WT^ cells, extending several microns in many cases (Fig. 5e). In contrast, MreB^WT^ formed small structures, presumably consisting of short filaments, with sizes consistent with structures imaged using epifluorescence microscopy (Fig. 5e). MreB^R124C^, MreB^D83E^, and MreB^A174T^ cells displayed fluorescence patch sizes (Methods) intermediate between those of MreB^WT^ and MreB^E276D^ cells (Fig. 5f), suggesting that all mutations stabilized filaments compared to MreB^WT^. Neither MreB^E276D^ nor MreB^WT^ structure sizes were significantly smaller when *rodZ* was deleted (Fig. 5f), suggesting that RodZ does not alter MreB oligomerization. MreB^WT^+MreB^E276D*^ cells contained MreB structures of intermediate size (Fig. 5g), consistent with the hypothesis that MreB^WT^ and MreB^E276D^ subunits form hybrid filaments. Consistent with previous observations, cells that were more rod-like (Methods) had larger MreB structures than did less-rod-like cells (Fig. S10, *p* < 0.0001 by the two-sample Kolmogorov-Smirnov test), suggesting that intermediate polymer size is required for the recovery of rod-shaped cells in the absence of RodZ.

## Discussion

Here, we demonstrate that *E. coli* dynamically modulates the geometric sensing of MreB via RodZ to drive changes in cell shape. Increased RodZ expression systematically enhanced the enrichment of MreB in regions of negative contour curvature (Fig. 2d), suggesting that RodZ alters the biophysical properties of MreB filaments, and decreased cell width (Fig. 2e), indicating that the changes in MreB localization affect cell morphology. Our MD simulations predicted that the bending of ATP-bound MreB filaments is reduced by RodZ binding (Fig. 4d), which could stabilize filaments on the relatively flat membrane. Nonetheless, it is also likely that filament mechanics is intrinsically coupled to biochemical parameters such as hydrolysis state [21], which in turn affect polymer size. Although structured illumination microscopy suggested that the formation of larger MreB oligomers with RodZ than without RodZ is necessary for rod-shaped growth (Fig. 5f), the observation that MreB^E276D^ did not rescue rod-like shape in Δ*rodZ* cells (Fig. S5d) may be due to the inability of very long filaments (Fig. 5g) to adjust to local variations in cell shape. By contrast, the combination of MreB^WT^ and MreB^E276D^ was sufficient to recover rod-shaped growth in the absence of RodZ (Fig. 5a-d), implying the need for balance between the polymeric properties of RodZ-bound and unbound MreB^WT^ filaments. The three Δ*rodZ* suppressor MreB mutants that we studied have different curvature enrichment profiles (Fig. 3d) and larger polymer patch sizes (Fig. 5f) than wild-type cells *in vivo,* and while these changes enable growth in the absence of Δ*rodZ*, potentially due to the capacity of these mutant filaments to mimic the effects of RodZ binding on MreB polymer mechanics (Fig. 4d), all three mutants have a fitness cost and altered shape relative to MreB^WT^ that we suggest results from the inability to properly modulate MreB filament length and mechanics in a wild-type manner. A previous study reported that the only Δ*rodZ* suppressor MreB mutant that had higher self-interaction than MreB^WT^ was also the only mutant that rescued rod-like shape in minimal medium [30], possibly due to the need for longer MreB filaments in minimal medium, further highlighting the links between regulation of MreB polymerization and shape determination. Inducible RodZ expression and MreB mutations should prove powerful and complementary tuning knobs for further dissection of the variables dictating MreB localization and cell-shape determination.

Much remains to be discovered regarding the links between MreB, its binding partners, and cell-wall insertion. A previous study showed that RodZ is required for processive motion of MreB [36], while our previous simulations suggested that curvature-mediated patterning could be responsible for processive motion along the circumferential deformations of negative Gaussian curvature induced by cell-wall insertion [11]. Given that RodZ also affects MreB curvature enrichment, the two bases for processivity are not necessarily contradictory. Moreover, while MreB depolymerization by A22 alters the pattern of cell-wall insertion, suggesting that MreB patterning dictates cell shape, it may also be the case that other aspects of cell size changes affect MreB dynamics and localization. In organisms such as *Bacillus subtilis* that have multiple MreB homologs, it is possible that RodZ differentially modulates the curvature enrichment of each homolog. MreB has been shown to colocalize with MreC/D [16] and FtsZ in *E. coli* [37]; in the latter case, FtsZ adopts various filament conformations [38, 39], which could couple mechanically to MreB. Thus, MreB may have as diverse a set of partners as the actin-binding proteins that enable myriad functions [40]. Given that some actin-binding proteins can deform membranes [41], other bacterial proteins may act similarly to RodZ to specifically modulate MreB’s curvature preference.

The rapid changes in cell shape during the progression from stationary phase to exponential growth and back (Fig. 1b) are consistent with the classic Growth Law of a positive relationship between nutrient-determined steady-state growth rate and cell size [23], as well as with a more recent finding linking relative rates of surface and volume synthesis to cell-size determination [42]. Our discovery that the curvature preference of MreB (Fig. 1h) varies continuously with growth rate (Fig. 1a), cell size (Fig. 1b), and the ratio of MreB to RodZ (Fig. 2d) suggests that RodZ-driven MreB localization may be a major component of the regulation of cell size; MreB enrichment profiles and cellular dimensions both changed gradually over the first 3 h of passaging (Fig. 1b,h), and the variability in enrichment profiles across time points was similar to what we achieved by modulating RodZ levels (Fig. 2d). Our data indicate that for a fixed contour curvature, MreB prefers wider regions of the cell (Fig. 1f), which may provide a homeostasis mechanism for cell width. The rapid dynamics in mean width and length over 1-2 h (Fig. 1b) indicate that both dimensions can be tuned over a few generations in either direction. This rapid size variation should be useful for probing many general physiological questions such as the coupling between DNA replication and cell division [43]. The remarkable ability of bacterial cells to adjust their size, and hence their physiology, with different concentrations of the same molecular components highlights their ability to regulate and exploit the biophysics of their cytoskeletons.

## Methods

### Strains and growth conditions

All *E. coli* strains and plasmids used in this study, along with the condition-dependent mean cell length and width of all imaging experiments, are described in Table S2. Strain construction was performed using standard transformation or transduction methods. Lysogeny broth (LB) with 5 g/L NaCl was used for all experiments. Strains were grown at 37 °C. The antibiotics chloramphenicol (Sigma-Aldrich) and cephalexin (MP Biomedicals) were used at concentrations of 15 μg/mL and 10 μg/mL, respectively. For *rodZ* induction experiments, xylose or arabinose were supplemented as described in the main text.

Δ*rodZ* suppressor MreB mutants were generated using λ-Red recombination in the parental strain expressing a sandwich fusion of MreB to msfGFP (NO34) following standard protocols [44]. The resulting colonies were confirmed by colony PCR and sequencing. Mutated MreB-msfGFP alleles were moved to a clean MG1655 background using P1 transduction.

To measure growth curves, cells were cultured in LB to stationary phase for 18 h or 24 h, then back diluted 200-fold into LB. Optical density was measured using an M200 plate reader (Tecan Group).

### Morphological time course

To monitor cell shape as a function of population density, we back-diluted a 24-h culture grown in LB 1:200 into fresh LB in a test tube. Every 15 min, we extracted a small sample and imaged cells on an agarose pad. To minimize temperature fluctuations of the growing culture, cultures were immediately returned to the incubator after the brief period of sample extraction. Cells were imaged with phase-contrast and epifluorescence microscopy to measure cell shape and MreB localization.

### Microscopy

Cells were imaged on a Nikon Eclipse Ti-E inverted fluorescence microscope with a 100X (NA 1.40) oil-immersion objective (Nikon Instruments Inc., Melville, NY, USA). Images were collected using an Andor DU885 EMCCD or Neo 5.5 sCMOS camera (Andor Technology, South Windsor, CT, USA). Cells were maintained at 37 °C during imaging with an active-control environmental chamber (HaisonTech, Taipei, Taiwan). Images were collected using μManager v. 1.4 [45].

### Image analysis

The MATLAB (MathWorks, Natick, MA, USA) image processing software *Morphometrics* [29] was used to segment cells and to identify cell contours from phase-contrast images. Fluorescence intensity profiles were generated by integrating image fluorescence along lines perpendicular to the contour at points uniformly spaced by approximately one pixel, extending five pixels in either direction. Mid-plane contour curvature was a three-point measurement defined by the arc-length derivative of the vector field formed from the unit normals to the contour, and did not assume any correlation of curvature values on opposite sides of the neutral axis of the cell [11]. Each curvature profile was smoothed with a low-pass Gaussian filter. Cells were categorized as rod-like or non-rod-like based on the success or failure of the *Morphometrics* meshing function, which determines whether a grid of lines perpendicular to a midline can be used as a coordinate system for the polygon defined by the cell contour.

### Equilibrium MD simulations

All simulations were performed as in Ref. [21] using the package NAMD [46] with the CHARMM27 force field [47, 48]. Water molecules were described with the TIP3P model [49]. Long-range electrostatic forces were evaluated with the particle-mesh Ewald summation approach with a grid spacing of <1 Å. An integration time step of 2 fs was used [50]. Bonded terms and short-range, nonbonded terms were evaluated at every time step, and long-range electrostatics were evaluated at every other time step. Constant temperature (*T* = 310 K) was maintained using Langevin dynamics, with a damping coefficient of 1.0 ps^-1^. A constant pressure of 1 atm was enforced using the Langevin piston algorithm [51] with a decay period of 200 fs and a time constant of 50 fs.

### Simulated systems

Twelve MD systems were analyzed (Table S3), eight of which were simulated in this study. For all simulations without RodZ, the MreB crystal structure of *T. maritima* MreB bound to AMP-PNP (PDB ID: 1JCG) [52] was used, with the nucleotide modified to ATP or ADP [21]. For simulations including RodZ, the cytoplasmic tail of RodZ was added to each MreB by aligning the simulated dimer with the co-crystal structure of MreB and RodZ (PDB ID: 2WUS). Water and neutralizing ions were added around each MreB dimer, resulting in final simulation sizes of 95,000-143,000 atoms. All unconstrained simulations were performed for at least 50 ns. Setup, analysis, and rendering of the simulation systems were performed with VMD [53]. To compute average values and distributions of measurements, only the last 30 ns of each simulation trajectory were used. To ensure that the simulations had reached equilibrium, measurement distributions were fit to a Gaussian distribution. A satisfactory fit implies that the system is located within an energy minimum well approximated by a harmonic potential. All simulations were repeated at least twice, and repeat simulations gave similar results (Fig. S7). Relative bending orientations of dimer subunits were calculated by determining the rotational transformation required to align the subunits [21].

### Sample preparation and imaging for structured illumination microscopy

Overnight, saturated cultures were back-diluted 1:100 into fresh LB and grown at 37 °C with shaking until 0D~0.1. One milliliter of the cells was fixed in phosphate-buffered saline containing 3% glutaraldehyde/3% paraformaldehyde (Electron Microscopy Sciences) at room temperature for 15 min, with 1 μg/mL FM4-64FX membrane stain (Invitrogen) added during fixation. Cells were washed three times in cold phosphate-buffered saline, and 1 μL of the cell solution was pipetted onto a No. 1.5 coverslip (Zeiss) coated with poly-L-lysine solution (Sigma-Aldrich). After the droplet dried, a small drop of ProLong Diamond AntiFade Mountant (Thermo Fisher) was added on top of the droplet, and the coverslip was mounted on a glass slide (VWR) and sealed with VALAP (equal parts Vaseline, lanolin, and paraffin by weight).

Cell samples were imaged on an OMX V4 microscope platform (GE Life Sciences) with a 100X (NA 1.42) oil-immersion objective (Nikon Instruments). Images from two channels were collected on two Evolve 512 electron-multiplying charged couple device cameras (Photometrics) using DeltaVision microscopy imaging system v. 3.70 (GE Life Sciences). The two-sample Kolmogorov-Smirnov test tests the null hypothesis that two one-dimensional samples are drawn from the same underlying probability distribution.

### Image analysis for structured illumination microscopy

Raw images were reconstructed and aligned using SoftWoRx v6.5.2 (GE Life Sciences), and maximum projection images were created using FIJI [54]. Individual cells were segmented by the FM4-64FX signal using *Morphometrics*. MreB patches within each cell contour were identified from the GFP channel based on intensity, and patches smaller than the diffraction limit for structured illumination microscopy (~0.03 μm^2^) were excluded from further quantification.

### Code/data availability

The datasets generated and/or analyzed during the current study and analysis software are available from the corresponding author on reasonable request.

## Author Contributions

A.C. and K.C.H. conceptualized the study. A.C., H.S., and K.C.H. designed the experiments. A.C. and H.S. performed cloning and single-cell imaging. A.C. performed molecular dynamics simulations. A.C., H.S., and K.C.H. analyzed data. A.C., H.S., and K.C.H. wrote the manuscript. All authors reviewed the manuscript before submission.

## Acknowledgments

The authors thank the Huang lab for helpful discussions, Piet de Boer, Felipe Bendezu, Nickolay Ouzounov, Joshua Shaevitz, and Zemer Gitai for strains, and the Stanford University Mass Spectrometry facility for assistance. This work was supported in part by a Stanford Graduate Fellowship and a Gerald J. Lieberman Fellowship (to A.C.), an Agilent Graduate Fellowship and a Stanford Interdisciplinary Graduate Fellowship (to H.S.), and NSF CAREER Award MCB-1149328, NIH Director’s New Innovator Award DP2-OD006466, the Allen Discovery Center at Stanford University on Systems Modeling of Infection, and the Stanford Center for Systems Biology under NIH grant P50-GM107615 (to K.C.H.). Structured illumination microscopy in this study was supported, in part, by Award Number 1S10OD01227601 from the National Center for Research Resources. The contents of this study are solely the responsibility of the authors and do not necessarily represent the official views of the National Center for Research Resources or the National Institutes of Health.

## References

1. Scheffers, D.-J., and Pinho, M.G. (2005). Bacterial cell wall synthesis: new insights from localization studies. Microbiology and molecular biology reviews: MMBR 69, 585–607.

2. Höltje, J.V. (1998). Growth of the stress-bearing and shape-maintaining murein sacculus of Escherichia coli. Microbiology and molecular biology reviews: MMBR 62, 181–203.

3. Tropini, C., Lee, T.K., Hsin, J., Desmarais, S.M., Ursell, T., Monds, R.D., and Huang, K.C. (2014). Principles of bacterial cell-size determination revealed by cell-wall synthesis perturbations. Cell reports 9, 1–8.

4. Wang, S., Furchtgott, L., Huang, K.C., and Shaevitz, J.W. (2012). Helical insertion of peptidoglycan produces chiral ordering of the bacterial cell wall. Proceedings of the National Academy of Sciences of the United States of America 109, E595–604.

5. van den Ent, F., Amos, L.a., and Löwe, J. (2001). Prokaryotic origin of the actin cytoskeleton. Nature 413, 39–44.

6. van Teeffelen, S., Wang, S., Furchtgott, L., Huang, K.C., Wingreen, N.S., Shaevitz, J.W., and Gitai, Z. (2011). The bacterial actin MreB rotates, and rotation depends on cell-wall assembly. Proceedings of the National Academy of Sciences of the United States of America 108, 15822–15827.

7. Salje, J., van den Ent, F., de Boer, P., and Löwe, J. (2011). Direct membrane binding by bacterial actin MreB. Molecular cell 43, 478–487.

8. Lee, T.K., Tropini, C., Hsin, J., Desmarais, S.M., Ursell, T.S., Gong, E., Gitai, Z., Monds, R.D., and Huang, K.C. (2014). A dynamically assembled cell wall synthesis machinery buffers cell growth. Proceedings of the National Academy of Sciences of the United States of America 111, 4554–4559.

9. Domínguez-Escobar, J., Chastanet, A., Crevenna, A.H., Fromion, V., Wedlich-Söldner, R., and Carballido-López, R. (2011). Processive movement of MreB-associated cell wall biosynthetic complexes in bacteria. Science (New York, N.Y.) 333, 225–228.

10. Garner, E.C., Bernard, R., Wang, W., Zhuang, X., Rudner, D.Z., and Mitchison, T. (2011). Coupled, circumferential motions of the cell wall synthesis machinery and MreB filaments in B. subtilis. Science (New York, N.Y.) 333, 222–225.

11. Ursell, T.S., Nguyen, J., Monds, R.D., Colavin, A., Billings, G., Ouzounov, N., Gitai, Z., Shaevitz, J.W., and Huang, K.C. (2014). Rod-like bacterial shape is maintained by feedback between cell curvature and cytoskeletal localization. Proceedings of the National Academy of Sciences of the United States of America.

12. Dye, N.A., Pincus, Z., Fisher, I.C., Shapiro, L., and Theriot, J.A. (2011). Mutations in the nucleotide binding pocket of MreB can alter cell curvature and polar morphology in Caulobacter. Molecular microbiology 81, 368–394.

13. Kruse, T., Møller-Jensen, J., Løbner-Olesen, A., and Gerdes, K. (2003). Dysfunctional MreB inhibits chromosome segregation in Escherichia coli. The EMBO journal 22, 5283–5292.

14. Monds, R.D., Lee, T.K., Colavin, A., Ursell, T., Quan, S., Cooper, T.F., and Huang, K.C. (2014). Systematic Perturbation of Cytoskeletal Function Reveals a Linear Scaling Relationship between Cell Geometry and Fitness. Cell reports 9, 1528–1537.

15. Ouzounov, N., Nguyen, J.P., Bratton, B.P., Jacobowitz, D., Gitai, Z., and Shaevitz, J.W. (2016). MreB Orientation Correlates with Cell Diameter in Escherichia coli. Biophysical Journal 111, 1035–1043.

16. Kruse, T., Bork-Jensen, J., and Gerdes, K. (2005). The morphogenetic MreBCD proteins of Escherichia coli form an essential membrane-bound complex. Molecular microbiology 55, 78–89.

17. Bendezú, F.O., Hale, C.a., Bernhardt, T.G., and de Boer, P.a.J. (2009). RodZ (YfgA) is required for proper assembly of the MreB actin cytoskeleton and cell shape in E. coli. The EMBO journal 28, 193–204.

18. Bean, G.J., Flickinger, S.T., Westler, W.M., McCully, M.E., Sept, D., Weibel, D.B., and Amann, K.J. (2009). A22 disrupts the bacterial actin cytoskeleton by directly binding and inducing a low-affinity state in MreB. Biochemistry 48, 4852–4857.

19. Gitai, Z., Dye, N.A., Reisenauer, A., Wachi, M., and Shapiro, L. (2005). MreB actin-mediated segregation of a specific region of a bacterial chromosome. Cell 120, 329–341.

20. Billings, G., Ouzounov, N., Ursell, T., Desmarais, S.M., Shaevitz, J., Gitai, Z., and Huang, K.C. (2014). De novo morphogenesis in L-forms via geometric control of cell growth. Molecular microbiology, 1–14.

21. Colavin, A., Hsin, J., and Huang, K.C. (2014). Effects of polymerization and nucleotide identity on the conformational dynamics of the bacterial actin homolog MreB. Proceedings of the National Academy of Sciences of the United States of America 111, 3585–3590.

22. Sezonov, G., Joseleau-Petit, D., and D’Ari, R. (2007). Escherichia coli physiology in Luria-Bertani broth. Journal of bacteriology 189, 8746–8749.

23. Schaechter, M., MaalOe, O., and Kjeldgaard, N.O. (1958). Dependency on Medium and Temperature of Cell Size and Chemical Composition during Balanced Growth of Salmonella typhimurium. Journal of General Microbiology 19, 592–606.

24. Peters, J.M., Colavin, A., Shi, H., Czarny, T.L., Larson, M.H., Wong, S., Hawkins, J.S., Lu, C.H., Koo, B.M., Marta, E., et al. (2016). A Comprehensive, CRISPR-based Functional Analysis of Essential Genes in Bacteria. Cell 165, 1493–1506.

25. Auer, G.K., Lee, T.K., Rajendram, M., Cesar, S., Miguel, A., Huang, K.C., and Weibel, D.B. (2016). Mechanical Genomics Identifies Diverse Modulators of Bacterial Cell Stiffness. Cell systems 2, 402–411.

26. Baba, T., Ara, T., Hasegawa, M., Takai, Y., Okumura, Y., Baba, M., Datsenko, K.A., Tomita, M., Wanner, B.L., and Mori, H. (2006). Construction of Escherichia coli K-12 in-frame, single-gene knockout mutants: the Keio collection. Molecular systems biology 2, 2006 0008.

27. Alyahya, S.A., Alexander, R., Costa, T., Henriques, A.O., Emonet, T., and Jacobs-Wagner, C. (2009). RodZ, a component of the bacterial core morphogenic apparatus. Proceedings of the National Academy of Sciences of the United States of America 106, 1239–1244.

28. Shiomi, D., Sakai, M., and Niki, H. (2008). Determination of bacterial rod shape by a novel cytoskeletal membrane protein. The EMBO journal 27, 3081–3091.

29. Ursell, T., Lee, T.K., Shiomi, D., Shi, H., Tropini, C., Monds, R.D., Colavin, A., Billings, G., Bhaya-Grossman, I., Broxton, M., et al. (2016). Rapid, precise quantification of bacterial cellular dimensions across a genomic-scale knockout library.

30. Shiomi, D., Toyoda, A., Aizu, T., Ejima, F., Fujiyama, A., Shini, T., Kohara, Y., and Niki, H. (2013). Mutations in cell elongation genes mreB, mrdA and mrdB suppress the shape defect of RodZ-deficient cells. Molecular microbiology 87, 1029–1044.

31. Li, G.W., Burkhardt, D., Gross, C., and Weissman, J.S. (2014). Quantifying absolute protein synthesis rates reveals principles underlying allocation of cellular resources. Cell 157, 624–635.

32. Schmidt, A., Kochanowski, K., Vedelaar, S., Ahrne, E., Volkmer, B., Callipo, L., Knoops, K., Bauer, M., Aebersold, R., and Heinemann, M. (2015). The quantitative and condition-dependent Escherichia coli proteome. Nature biotechnology 34, 104–110.

33. Van den Ent, F., Izoré, T., Bharat, T.A., Johnson, C.M., and Löwe, J. (2014). Bacterial actin MreB forms antiparallel double filaments. eLife, e02634.

34. de Pedro, M.A., Young, K.D., Holtje, J.V., and Schwarz, H. (2003). Branching of Escherichia coli cells arises from multiple sites of inert peptidoglycan. Journal of bacteriology 185, 1147–1152.

35. Hedge, P.J., and Spratt, B.G. (1985). Amino acid substitutions that reduce the affinity of penicillin-binding protein 3 of Escherichia coli for cephalexin. European journal of biochemistry / FEBS 151, 111–121.

36. Morgenstein, R.M., Bratton, B.P., Nguyen, J.P., Ouzounov, N., Shaevitz, J.W., and Gitai, Z. (2015). RodZ links MreB to cell wall synthesis to mediate MreB rotation and robust morphogenesis. Proceedings of the National Academy of Sciences of the United States of America 112, 12510–12515.

37. Fenton, A.K., and Gerdes, K. (2013). Direct interaction of FtsZ and MreB is required for septum synthesis and cell division in Escherichia coli. The EMBO journal 32, 1953–1965.

38. Hsin, J., Gopinathan, A., and Huang, K.C. (2012). Nucleotide-dependent conformations of FtsZ dimers and force generation observed through molecular dynamics simulations. Proceedings of the National Academy of Sciences of the United States of America 109, 9432–9437.

39. Li, Y., Hsin, J., Zhao, L., Cheng, Y., Shang, W., Huang, K.C., Wang, H.W., and Ye, S. (2013). FtsZ protofilaments use a hinge-opening mechanism for constrictive force generation. Science 341, 392–395.

40. Winder, S.J., and Ayscough, K.R. (2005). Actin-binding proteins. Journal of cell science 118, 651–654.

41. Itoh, T., Erdmann, K.S., Roux, A., Habermann, B., Werner, H., and De Camilli, P. (2005). Dynamin and the actin cytoskeleton cooperatively regulate plasma membrane invagination by BAR and F-BAR proteins. Developmental cell 9, 791–804.

42. Harris, L.K., and Theriot, J.A. (2016). Relative Rates of Surface and Volume Synthesis Set Bacterial Cell Size. Cell 165, 1479–1492.

43. Wallden, M., Fange, D., Lundius, E.G., Baltekin, O., and Elf, J. (2016). The Synchronization of Replication and Division Cycles in Individual E. coli Cells. Cell 166, 729–739.

44. Sharan, S.K., Thomason, L.C., Kuznetsov, S.G., and Court, D.L. (2009). Recombineering: a homologous recombination-based method of genetic engineering. Nat Protoc 4, 206–223.

45. Edelstein, A., Amodaj, N., Hoover, K., Vale, R., and Stuurman, N. (2010). Computer control of microscopes using microManager. Current protocols in molecular biology / edited by Frederick M. Ausubel … [et al.] *Chapter* 14, Unit14 20.

46. Phillips, J.C., Braun, R., Wang, W., Gumbart, J., Tajkhorshid, E., Villa, E., Chipot, C., Skeel, R.D., Kale, L., and Schulten, K. (2005). Scalable molecular dynamics with NAMD. Journal of computational chemistry 26, 1781–1802.

47. MacKerell, A.D., Bashford, D., Bellott, Dunbrack, R.L., Evanseck, J.D., Field, M.J., Fischer, S., Gao, J., Guo, H., Ha, S., et al. (1998). All-Atom Empirical Potential for Molecular Modeling and Dynamics Studies of Proteins. The Journal of Physical Chemistry B 102, 3586–3616.

48. Foloppe, N., and MacKerell, J.A.D. (2000). All-atom empirical force field for nucleic acids: I. Parameter optimization based on small molecule and condensed phase macromolecular target data. Journal of computational chemistry 21, 86–104.

49. Jorgensen, W.L., Chandrasekhar, J., Madura, J.D., Impey, R.W., and Klein, M.L. (1983). Comparison of simple potential functions for simulating liquid water. The Journal of Chemical Physics 79, 926–935.

50. Tuckerman, M., Berne, B.J., and Martyna, G.J. (1992). Reversible multiple time scale molecular dynamics. The Journal of Chemical Physics 97, 1990–2001.

51. Feller, S.E., Zhang, Y., Pastor, R.W., and Brooks, B.R. (1995). Constant pressure molecular dynamics simulation: The Langevin piston method. The Journal of Chemical Physics 103, 4613–4621.

52. van den Ent, F., Amos, L.A., and Lowe, J. (2001). Prokaryotic origin of the actin cytoskeleton. Nature 413, 39–44.

53. Humphrey, W., Dalke, A., and Schulten, K. (1996). VMD: visual molecular dynamics. Journal of molecular graphics 14, 33–38, 27-38.

54. Schindelin, J., Arganda-Carreras, I., Frise, E., Kaynig, V., Longair, M., Pietzsch, T., Preibisch, S., Rueden, C., Saalfeld, S., Schmid, B., et al. (2012). Fiji: an open-source platform for biological-image analysis. Nature methods 9, 676–682.

